# Mapping the unicellular transcriptome of the ascending thoracic aorta to changes in mechanosensing and mechanoadaptation during aging

**DOI:** 10.1101/2022.07.21.501037

**Authors:** Yasmeen M. Farra, Cristobal Rivera, Michele Silvestro, Jacqueline Matz, Yogi Pratama, Puja Kumari, John Vlahos, Bhama Ramkhelawon, Chiara Bellini

## Abstract

Aortic stiffening is an inevitable manifestation of chronological aging, yet the mechano-molecular programs that orchestrate region- and layer-specific adaptations along the length and through the wall of the aorta are incompletely defined. Here, we show that the decline in passive cyclic distensibility is more pronounced in the ascending thoracic (ATA) compared to distal segments of the aorta and that tissues in both the medial and adventitial compartments of the ATA stiffen during aging. Single-cell RNA sequencing of aged ATA tissues reveals altered cellular senescence, remodeling, and inflammatory responses accompanied by enrichment of T-lymphocytes and rarefaction of vascular smooth muscle cells, compared to young samples. T-lymphocytes accumulate in the adventitia and likely promote fibrosis, while activation of mechanosensitive piezo-1 enhances medial vasoconstriction. These results portray the immuno-mechanical aging of the ATA as a process that culminates in a stiffer conduit permissive to the accrual of multi-gerogenic signals priming to disease development.

## INTRODUCTION

Epidemiological evidence has established that cardiovascular disease (CVD) is a leading cause of death amongst adults 65 and older^1^ and advanced age is one of the dominant risk factors that fuels the development of CVD^2^. Concurrent clinical findings of elevated body mass index, hypertension, hyperlipidemia, and smoking further enhance the risk of acute cardiovascular events in the aging population, including myocardial infarction, ischemic stroke, and heart failure^3^. Considering that chronological aging cannot be reversed, dissection of age-responsive molecular and cellular pathways and their impact on the function of central arteries is crucial to mitigate the damaging effects of aging and intercept aortic disease via spatio-temporal pharmacological interventions.

Vascular function is essentially mechanical in nature, and mechanical metrics have emerged as reliable estimators of cardiovascular age. Amongst those, structural stiffening of elastic arteries is an independent predictor of all-causes and cardiovascular mortality in hypertensive patients, constituting a risk factor for CVD in hypertensive and normotensive populations alike^4–6^. A spatially-averaged measure of structural stiffness, pulse wave velocity (PWV), doubles between 20 and 80 years of age^7^, thereby elevating the workload on the heart and contributing to negative downstream effects on microvasculature and organs^8^. Enzymatic degradation, fragmentation, and calcification of medial elastic material^9^, deposition of collagen throughout the aortic wall^9–11^, and cross-linking of structural proteins due to non-enzymatic glycation^12,13^ synergistically contribute to the age-related structural stiffening of conduit arteries. Notwithstanding these insights, the broad spectrum of molecular and cellular adaptations that accompany a compromised aged vasculature has yet to be fully resolved^14^.

We determined that the structural stiffening of the mouse aorta occurs progressively with aging and heterogeneously along the artery, with the proximal ascending thoracic aorta (ATA) experiencing the largest distensibility decline. Single-cell RNA sequencing of ATA samples isolated from 12- and 84-week-old mice revealed that signaling pathways such as cellular senescence, inflammation, vascular remodeling, and endothelial dysfunction are overrepresented in aged mice. Furthermore, while progressive adventitial fibrosis may follow T-lymphocyte infiltration in ATA tissues, enhancement of the vasoconstrictive response due to Piezo-1 activation suggests a role for mechanosensing on the age-induced medial remodeling. These processes collectively culminate in a stiffer conduit with reduced ability to store elastic energy for diastolic blood flow augmentation. Our work thus provides direct evidence of intrinsic vascular maladaptations during aging and identifies Piezo-1 as one of the gerogenic culprits of pathological ECM remodeling.

## RESULTS

### Accelerated structural stiffening in the ascending portion of the aged aorta

To evaluate whether different segments of the aorta age synchronously, we characterized the biaxial mechanical behavior of the ascending thoracic (ATA), descending thoracic (DTA), suprarenal abdominal (SAA), and infrarenal abdominal (IAA) aorta of young (12-week-old) and aged (84-week-old) mice (Figure 1A). The passive pressure vs. diameter (P-d) response of the aorta shifted toward larger diameters with age, though this effect progressively faded when moving away from the heart (Figure 1B). More pronounced in proximal segments was also the change in the shape of the P-d curve, consistent with an earlier engagement of collagen fibers and suggestive of decreased ability to deform within the physiological range of pressures^15^ (Figure 1B). Indeed, the ATA experienced the largest decrease in distensibility, which remained significant but muted in the DTA, then subsided in the abdominal segments (Figure 1C). As our results demonstrate accelerated effects of aging on the ascending segment of the thoracic aorta, we will focus on this region for the rest of the work.

**Figure 1.**
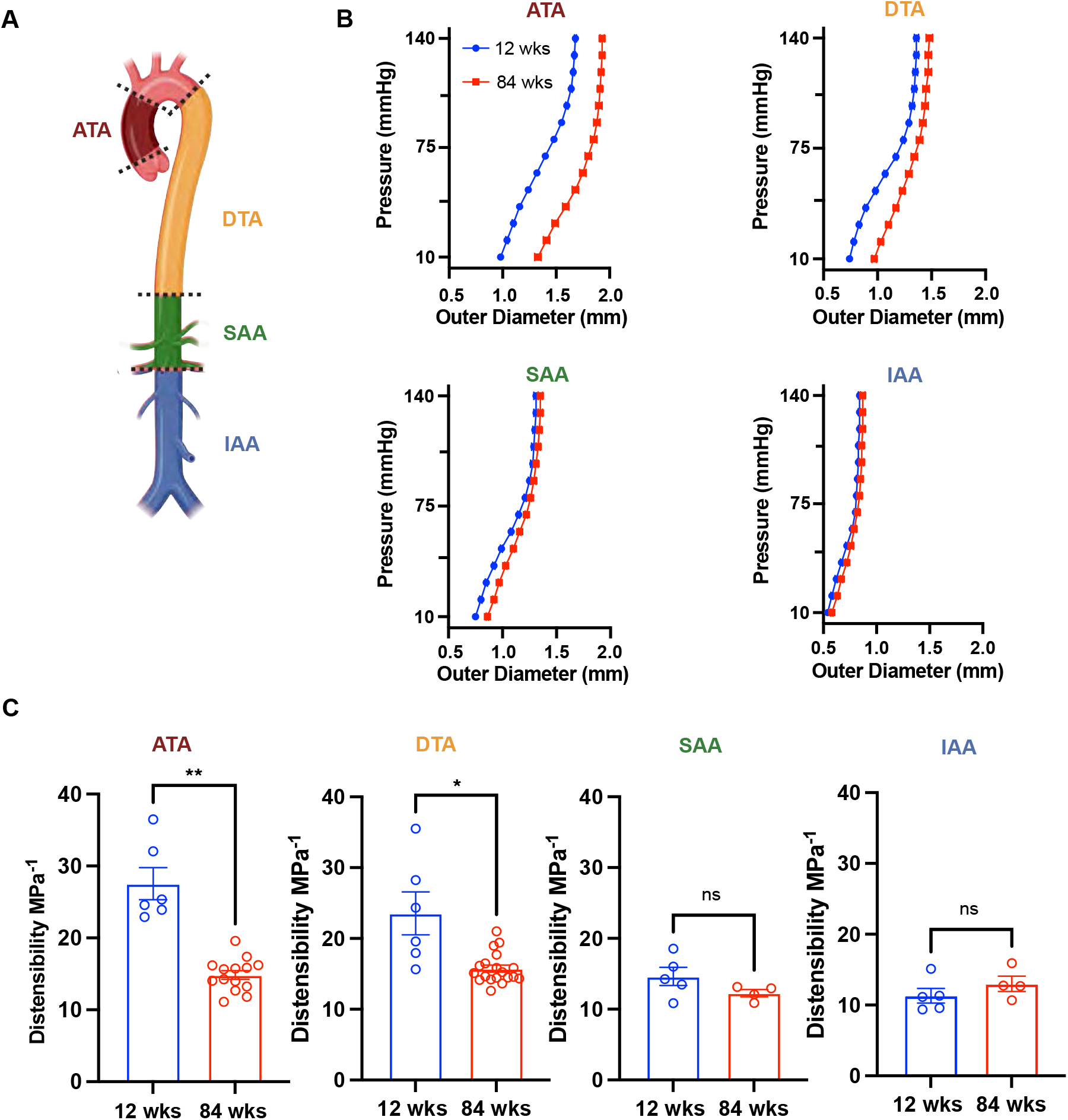
Spatial heterogeneity in the structural stiffening of the female murine thoracic aorta during aging. (A) Schematic detailing the location of the four anatomical segments of interest from proximal to distal: ascending (ATA) and descending (DTA) thoracic, and suprarenal (SAA) and infrarenal (IAA) abdominal aorta. (B) Passive pressure vs. outer diameter response at 12 (N=6) and 84 (N=14) weeks of age in the four aortic regions. (C) Cyclic aortic distensibility in thoracic and abdominal segments at the 12 and the 84-week endpoints. Statistical difference between age groups denoted by ** overbar for p < 0.01 and *** overbar for p < 0.001.

### Mapping unicellular transcriptome of the ascending thoracic aorta during aging

To unbiasedly catalogue the multicellular transcriptomic variations that arise during chronological aging of the ATA, we performed single-cell RNA sequencing (scRNA-seq) on tissues from 12- and 84-week-old mice, using the 10x Genomics platform. Three samples in each age group were digested and 15,000 cells from each of the 12- and 84-week-old ATAs were processed for integrated single-cell sequencing to detect the transcriptome (Figure 2A). Quality control consisted of removing cells that exhibited less than 5% of mitochondrial DNA. Unsupervised Seurat clustering of the single-cell data allowed identification of 12 aortic cell populations, as shown in the Uniform Manifold Approximation and Projection (UMAP) plots (Figure 2B). We utilized a canonical set of markers represented in the heatmap (Figure 2D) to define vascular cells – endothelial cells (*Pecam1, Cdh5*), vascular smooth muscle cells (vSMCs) (*Acta2, Myh11, and Myl9*), and aortic fibroblasts (*Col1a1, Col3a1, Dcn, Gsn*) (Figure 2C). Interestingly, there was an enrichment of immune cell populations characterized by T lymphocytes (*Cd3e, Cd8a, Trbc1, Trbc2*) in the aged ATA samples, and we observed a shift in the presence of vascular cells and T lymphocytes in the ATA with aging (Figure 2E). The proportion of T lymphocytes increased ∼30x in the aged ATA while the proportion of vSMC decreased ∼5x (Figure 2F-G). These results suggest that chronological aging impacts the qualitative and quantitative immuno-vascular phenotypes that populate the ATA.

**Figure 2.**
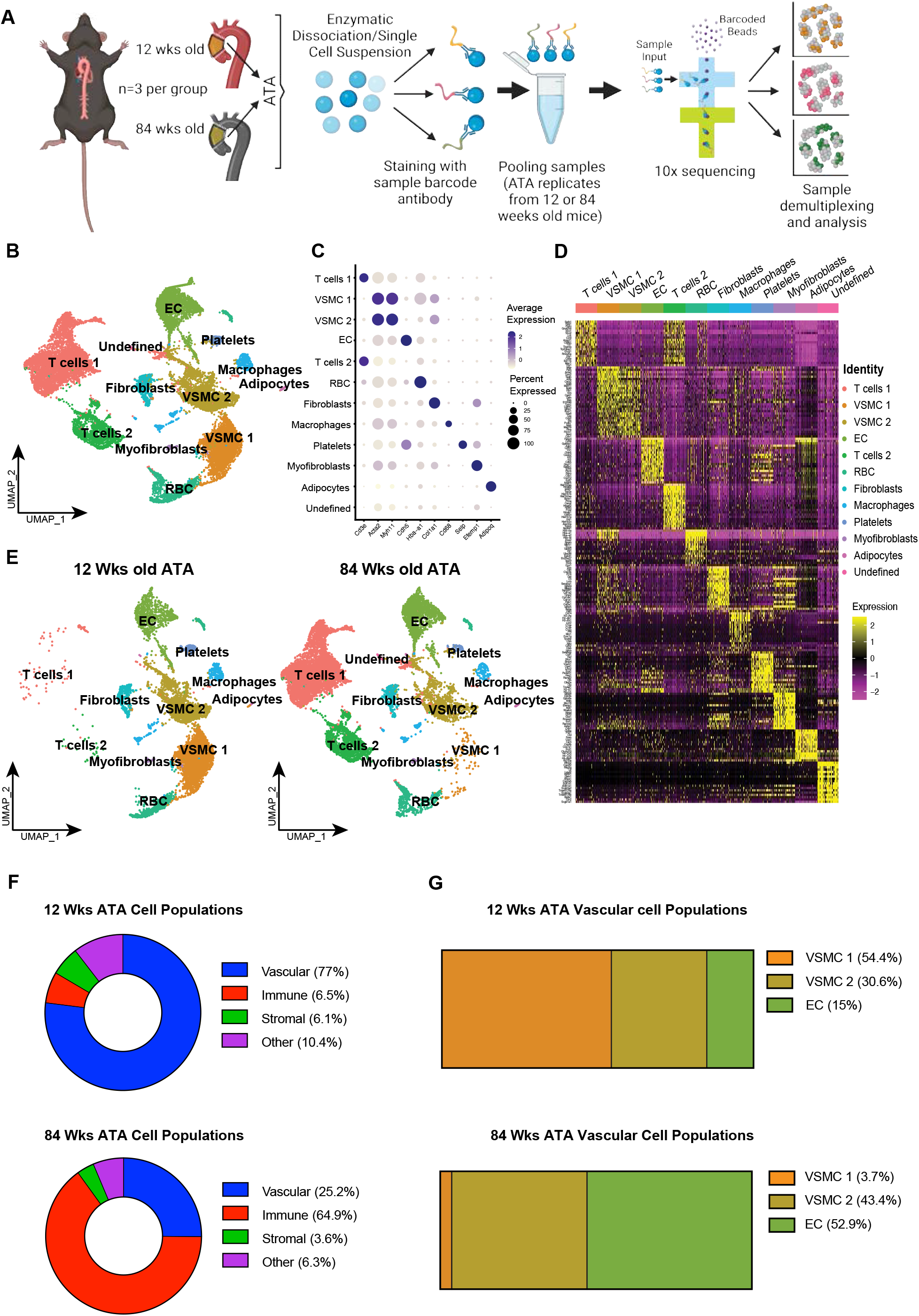
Mapping the unicellular transcriptome of the ATA during aging. (A) Schematic detailing sample collection, demultiplexing, and analysis. (B) Representative uniform manifold approximation and projection (UMAP) of cell clusters in the murine ATA. (C) Plot of cell clusters identified based on average expression of genes and the percent they are expressed. (D) Heatmap of top differentially expressed transcripts by each cell clusters using expression of canonical markers to tag vascular cells. (E) UMAP of ATA cell clusters at the 12- and 84-week endpoints. (F) Pie chart showing the relative makeup of 12- and 84-week-old cell populations based on category: vascular, immune, stromal, or other. (G) Bar chart visualizing relative quantities of vascular cell populations at 12 and 84 weeks.

### Diverse patterns of signaling pathways during aging of the ATA

To delve into the molecular pathways that are altered during aging, we performed enrichment pathway analysis using the Ingenuity Pathway Analysis (IPA) software. We observed an overrepresentation of cellular senescence, extracellular remodeling, and acquired immune response pathways in the ATA of 84-compared to 12-week-old mice (Figure 3A). In contrast, a decline in endothelial integrity, angiogenic responses, and autophagy was noted in aged ATAs compared to younger samples (Figure 3A). Expression of a battery of genes that regulate the autophagy machinery, including *Calm1, Gabarap, Lamp1, MAP1c3a* and *Map1lc3b* was significantly downregulated with age in our dataset (Figure 3E). Elevated cellular senescence in aged tissues was denoted by upregulation of a panel of markers previously reported in the literature^16–23^, namely *Ccnd3, Cdkn1b, Elf1, Pten, Ets2, Itpr2, Nfatc3*, and *Ets1* (Figure 3B). Furthermore, upregulated gene expression (Figure 3C) and enhanced immunofluorescent staining intensity (Figure 3D) confirmed extracellular accumulation of HMGB1 in aged vasculature compared to young ATA tissues, consistent with accrual of senescent molecular traits during aging (Figure 3B)^24^. These results indicate that several canonical pathways of aging are present in the ATA and are likely to contribute to pathological vascular remodeling.

**Figure 3.**
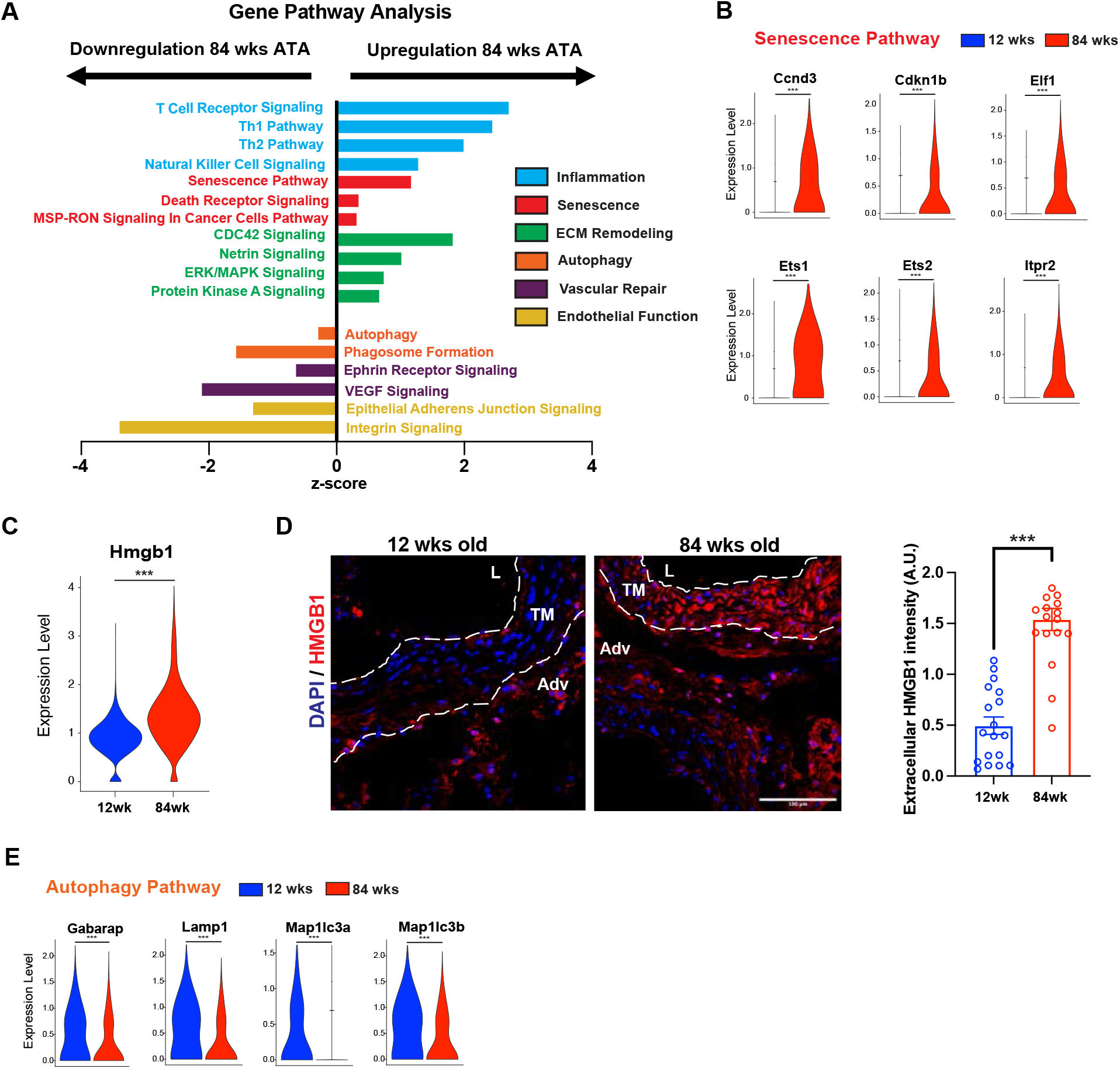
Diverse patterns of signaling pathways in the ATA during aging. (A) Up- and down-regulated signaling pathways in ATA tissues from 84-week-old mice, compared with the 12-week endpoint. (B) Multiple senescence pathway-associated genes in the aged ATA. (C) HMGB1 gene expression in 84-week-old tissues compared with the 12-week endpoint. (D) Immunofluorescent staining colocalized with DAPI (cell nuclei, blue) and HMGB1 (red) in the 84-week-old ATA, compared with 12-week-old tissues. (E) Multiple autophagy pathway-associated genes are downregulated in 84-week-old ATA samples. Statistical difference between age groups denoted by *** overbar for p < 0.001.

### Heterogeneous accumulation of T lymphocytes during aging of the ATA

Since T-cell receptor signaling is one of the most abundant signaling pathways in the aged ATA, we further dissected the nature of T lymphocyte clone distribution within this cluster. We identified distinct clusters of CD3d, e.g., CD4 and CD8 lymphocytes in our dataset (Figure 4A). Additional analysis of T-cell clones revealed an enrichment of naïve T lymphocytes in the aged aorta (Figure 4B). A similar trend was found in differentiated Th1 (Figure 4C), Th2 (Figure 4D), regulatory (Figure 4E), and Th17 (Figure 4F) T lymphocytes. We observed a surge of cytotoxic CD8 lymphocytes (Figure 4G) and elevated effector lymphocytes of the innate immune family (Figure 4H). Natural Killer (NK) cells expressing *Gzmb* (Granzyme B) and *Prf1* (Perforin1) transcripts (Figure 4I) were also detected. Furthermore, we noted an enrichment of the NK signaling pathway, characterized by expression of *Lat* (Linker for activation of T-cells family member 1) and *Lck* (lymphocyte-specific protein tyrosine kinase) by NK cells that populate the aged ATA (Figure 4J). Interestingly, we observed elevated expression of Rag1, a gene providing instructions for an essential component involved in the development and maturation of T and B cells^25^ (Figure 4L). Confirming transcriptomic increase of T lymphocyte signaling in the aged ATA, we measured elevated immunofluorescent CD3+ T-cells staining in the adventitia of ATA cross-sections at the 84-week endpoint (Figure 4M). Altogether, T lymphocytes receptor signaling was broadly elevated and activated in the aged ATA (Figure 4K), revealing a likely key role for the adaptive immune response in the remodeling of the aortic wall during aging.

**Figure 4.**
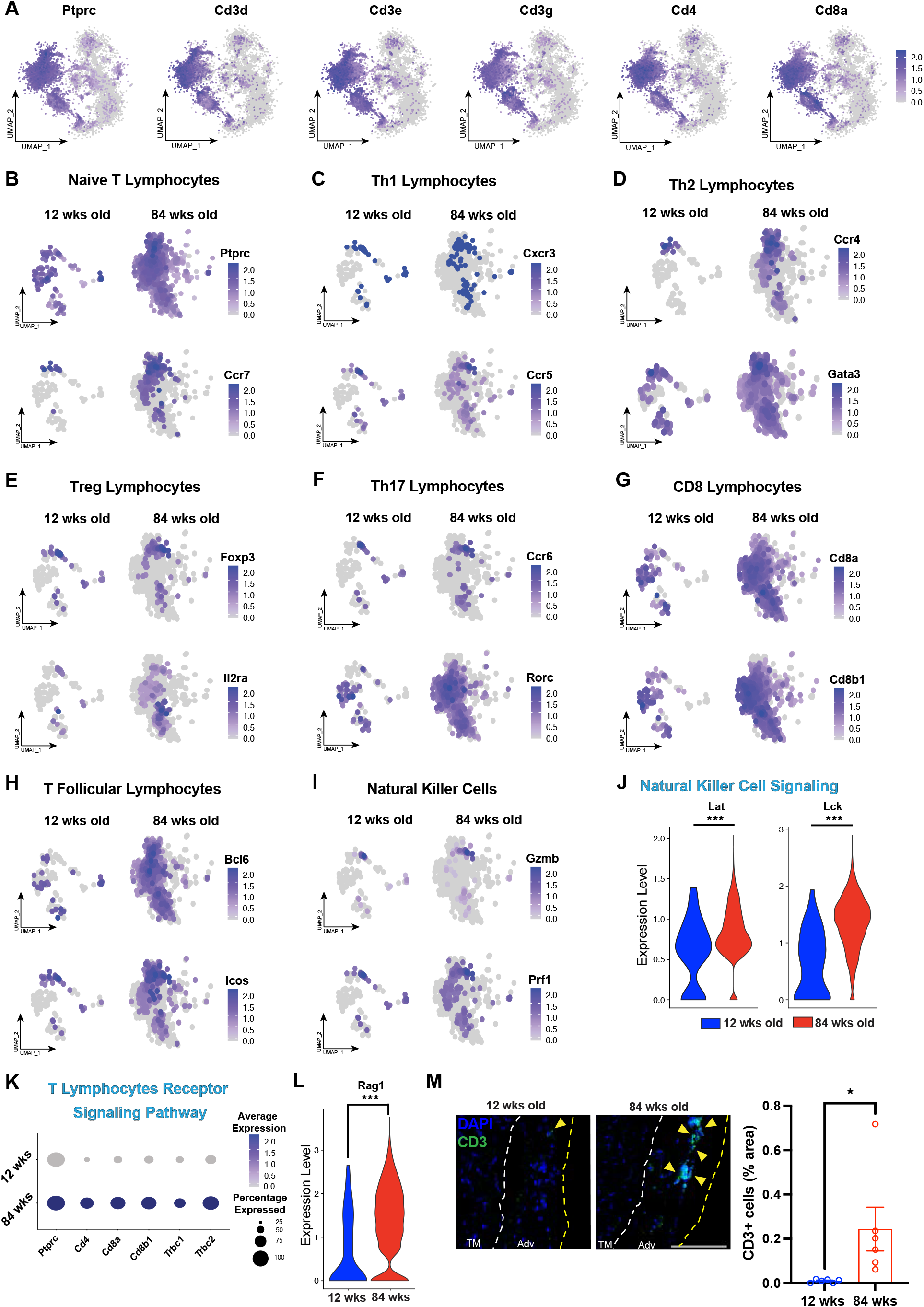
Heterogeneous accumulation of T lymphocytes in the ATA during aging. (A) ATA tissues feature distinct clusters of CD3d, e.g., CD4 and CD8 lymphocytes. Analysis of T-cell clones identified an enrichment of (B) naïve T lymphocytes and differentiated (C) Th1, (D) Th2, (E) regulatory, and (F) Th17 T lymphocytes in the aged ATA. The 84-week-old ATA exhibits (G) increased cytotoxic CD8 lymphocytes and (H) elevated effector lymphocytes of the innate immune family. (I) Natural Killer (NK) cells expressing *Gzmb* (Granzyme B) and *Prf1* (Perforin1) transcripts are detected in ATA tissues. (J) NK signaling pathway, characterized by expression of *Lat* (Linker for activation of T-cells family member 1) and *Lck* (lymphocyte-specific protein tyrosine kinase) by NK cells. (K) T lymphocyte receptor signaling in the ATA during aging. (L) Rag1 expression in the ATA at the 84-week endpoint. (M) Immunofluorescent staining colocalized with DAPI (cell nuclei, blue) for expression of CD3+ cells (CD3, green) in 84-week-old ATA tissues, compared to the 12-week endpoint, with Adv indicating adventitial area outlined between the white and yellow dotted lines, and TM representing the tunica media to the left of the white dotted line. Yellow arrows highlight the location of CD3+ cells. Scale bar is 100μm. Plot indicates relative fluorescent intensity. Statistical difference between age groups denoted by * overbar for p < 0.05 and *** overbar for p < 0.001.

### Microstructural matrix adaptations of the aortic wall during aging

Several markers comprising subfamilies that safeguard the extracellular matrix were altered transcriptionally in the aged ATA. While expression of elastin (*Eln*), decorin (*Dcn*), biglycan (*Bgn*), and tissue inhibitors of matrix metalloproteinases 2 and 3 (*Timp2, 3*) declined within the vSMC cluster of the aged ATA (Figure 5A), we noted upregulation of collagen superfamily elements in fibroblasts (Figure 5B). Consistently, *α*-smooth muscle actin expression in the adventitia was significantly larger in 84-vs. 12-week-old ATA samples (Figure 7A), suggesting that fibroblasts may differentiate into a synthetic, secretory phenotype with advancing age^26^. To corroborate transcriptional inferences, we determined the microstructural composition of the ATA wall at selected endpoints throughout the mouse adult lifespan (Figure S1) and found alterations in the overall makeup of tissue microstructure with age (Figure S2B). Histological analysis revealed that the area fraction of collagen in the adventitia broadened significantly between 12 and 49 weeks of age (67 ± 3% vs. 89 ± 5%, p = 0.016), after which it remained stable (Figure 5D). Similarly, the relative collagen content of the media expanded with age, but did so more gradually, rising from 9 ± 1% at 12 weeks to 36 ± 7% at 84 weeks (p = 0.01; Figure 5D). A significant positive correlation emerged between increasing age and the area fraction of collagen through the wall (Spearman’s correlation coefficient r_s_ = 0.643, p < 0.01; Figure S2A) and in each layer (r_s_ = 0.364, p < 0.01 for the adventitia and r_s_ = 0.692, p < 0.01 for the media; Figure 5D). As a result, the area fraction of elastin exhibited a significant negative correlation with age (r_s_ = -0.450, p < 0.01), despite no marked elastic fiber degradation (Figure 5C). Overall, these findings recognize collagen remodeling as a dominant feature of microstructural aging in the medial and adventitial layers of the ATA wall.

**Figure 5.**
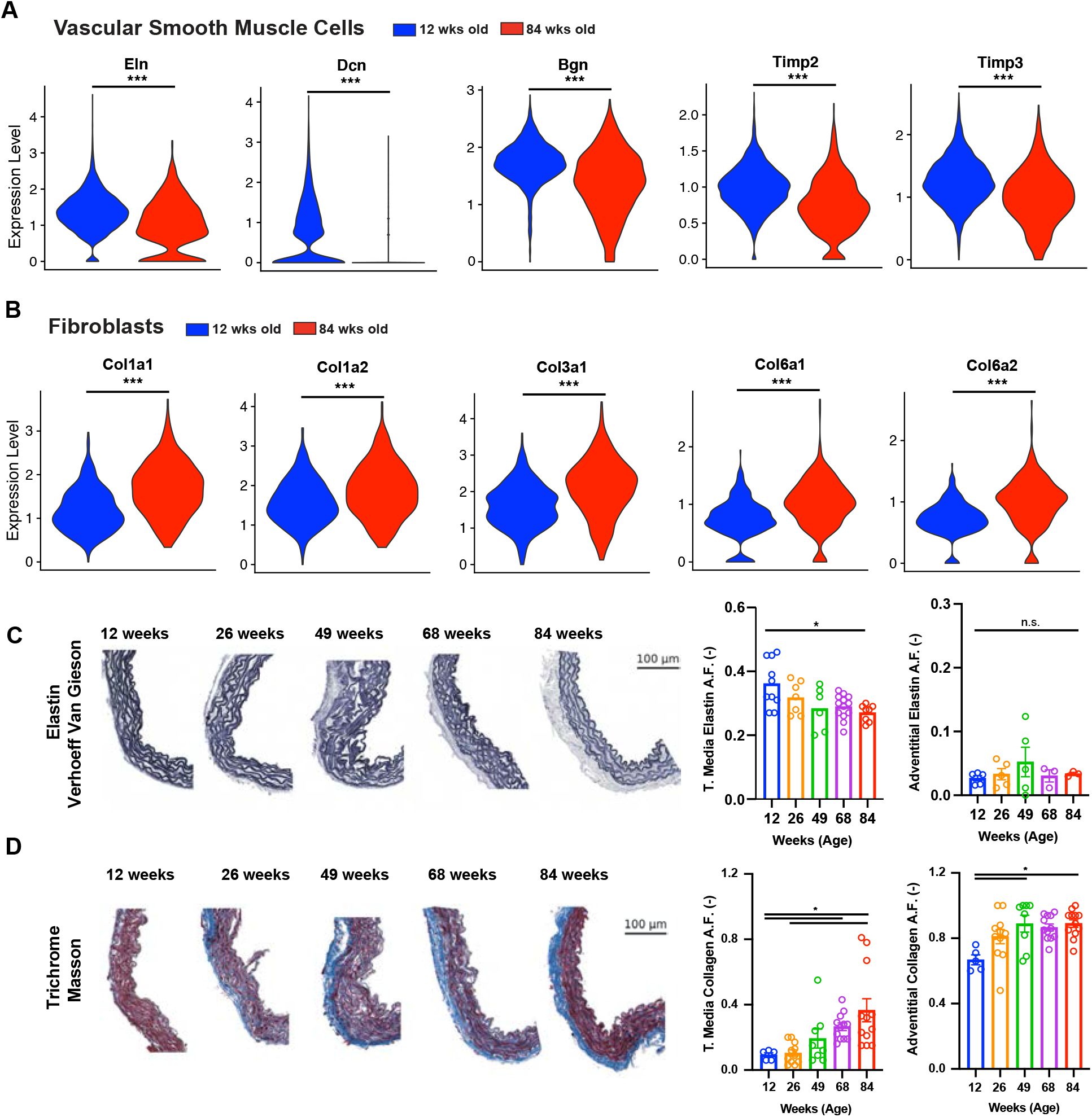
Microstructural matrix adaptations of ATA tissues during aging. (A) Expression of elastin (*Eln*), Decorin (*Dcn*), Biglycan (*Bgn*), and tissue inhibitors of matrix metalloproteinases 2 and 3 (*Timp2, 3*) genes in the vSMC cluster of the 12- and 84-week-old ATA. (B) Expression of collagen superfamily genes, including *Col1a1, Col1a2, Col3a1, Col6a1*, and *Col6a2* in ATA samples at 12 and 84 weeks of age. (C) Quantification of elastin content (black fibers) in the tunica media (T.M.) and adventitia of ATA cross sections stained with Verhoeff Van Gieson (VVG) stain at 12, 26, 49, 68, and 84 weeks of age. (D) Quantification of collagen (blue) and vSMC cytoplasm (red) content in the tunica media (T.M.) and adventitia of ATA cross sections stained with Masson’s Trichrome (MTC) stain at 12, 26, 49, 68, and 84 weeks of age. Statistical difference between age groups denoted by * overbar for p < 0.05 and *** overbar for p < 0.001.

### Shift toward dysfunctional passive aortic wall mechanics with aging

The remodeling endured by the ATA wall during aging may support a progressive shift toward a collagen-driven mechanical response, consistent with the large and significant decline in distensibility that we observed between 12 and 84 weeks of age (Figure 1C). To establish a timeline for this process, we performed inflation/extension tests on ATA samples at the same endpoints considered for microstructural analysis (Figure S1). Note, mice steadily gained weight until 68 weeks, when their body mass reached a plateau (Table S1). Importantly, systolic and diastolic values of peripheral blood pressure remained constant throughout (Table S1).

The passive response of the ATA gradually drifted with age. From a structural standpoint, a leftward shift in the axial force vs. length behavior complemented the rightward shift in the pressure vs. diameter curve (Figure 6A). Comparison of average biaxial stress vs. stretch curves across groups further detailed the effect of aging on the intrinsic behavior of tissues, with progressive loss of circumferential distensibility and axial extensibility (Figure 6A). For completeness, Table S2 reports the best-fit constitutive descriptors of the bulk (i.e., averaged through thickness) mechanical response of ATA tissues and Table S3 includes related predictions of morphological and mechanical parameters. Finally, Table S4 lists the best-fit coefficients that characterize the layer-specific mechanical properties of ATA tissues.

**Figure 6.**
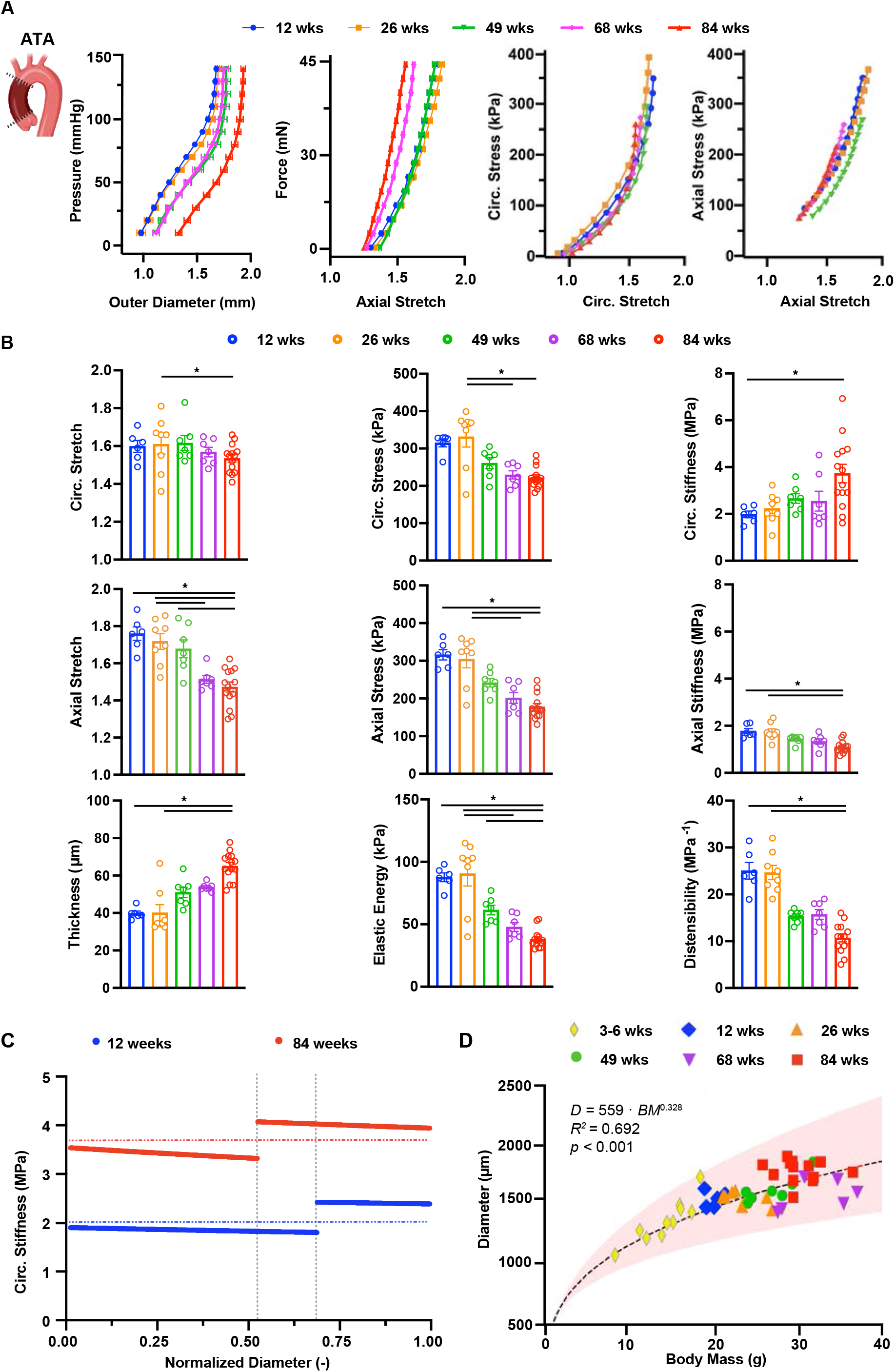
Progressive loss of mechanical functionality in the ATA during aging. (A) Average (± SEM) structural (pressure vs. outer diameter, axial force vs. stretch) and tissue (Cauchy stress vs. stretch) behavior of ATA samples in the circumferential and axial directions at 12, 26, 49, 68, and 84 weeks of age. (B) Biaxial descriptors of mechanical function in the ATA at the 5 endpoints. Metrics of stretch, stress, stiffness, thickness, and energy are calculated at group-specific values of systolic pressure. Cyclic distensibility accounts for group-specific values of luminal pressure and diameter between diastole and systole. (C) Linearized circumferential stiffness of tissues in the medial and adventitial layers of the 12- and 84-week-old ATA. Note, horizontal dash-dotted lines indicate through-thickness integral mean values of circumferential stiffness for the two groups, closely matching the predictions of the transmurally-averaged model (Table 3). Grey vertical dotted lines define the medial/adventitial border in the two groups. (D) Allometric scaling of systolic luminal diameter with body mass in ATA samples throughout the mouse lifespan. Group sizes for the passive mechanical analysis are: 12 weeks (N = 6), 26 weeks (N = 8), 49 weeks (N = 8), 68 weeks (N = 9), and 84 weeks (N = 19). Note, we incorporated data from developing female wildtype mice (N = 9) to improve allometric predictions at low body mass. Statistical significance between individual age groups denoted by * overbar for p < 0.05.

The aorta dilated as mice aged (r_s_ = 0.618, p < 0.01), with the unloaded luminal diameter expanding from 892 ± 43 μm at 12 weeks to 1060 ± 25 μm at 84 weeks. Likewise, the unloaded thickness of the aortic wall increased from 111 ± 1 μm at 12 weeks up to 146 ± 5 μm at 84 weeks (r_s_ = 0.808, p < 0.01). Thickening and luminal widening persisted when evaluated at physiological levels of pressure and axial stretch. Systolic luminal diameter rose from 1560 ± 24 μm at 12 weeks to 1782 ± 22 μm at 84 weeks (r_s_ = 0.632, p < 0.01), while systolic thickness increased from 40 ± 1 μm to 65 ± 2 μm between 12 and 84 weeks (r_s_ = 0.759, p < 0.01; Figure 6B). Because representative (through-thickness average) diameters when in the traction-free configuration and under the effect of physiological loads varied in a similar manner during aging, the circumferential stretch did not significantly correlate with age and only exhibited a significant difference between 26 and 84 weeks (1.61 ± 0.05 vs. 1.53 ± 0.02; Figure 6B). Since wall thickening outpaced the widening of the lumen, circumferential wall stress showed a negative correlation with age (r_s_ = -0.574, p < 0.01), whereby it declined after 26 weeks toward significantly lower values at 68 and 84 weeks (331 ± 27 kPa at 26 weeks vs. 222 ± 7 kPa at 84 weeks; Figure 6B). Nevertheless, the intrinsic stiffness of tissues in the circumferential direction increased with advancing age (r_s_ = 0.521, p = 0.001), from 1.98 ± 0.14 MPa at 12 weeks to 3.73 ± 0.40 MPa at 84 weeks (Figure 6B). Importantly, circumferential tissue stiffening between 12 and 84 weeks of age occurred in both the media and adventitial layers of the wall (Figure 6C).

The axial stretch steadily decreased with age (r_s_ = -0.738, p < 0.01), first becoming statistically different between 26 and 68 weeks (1.72 ± 0.04 vs. 1.51 ± 0.02; Figure 6B). Likewise, both axial stress (r_s_ = -0.707, p < 0.01) and axial tissue stiffness (r_s_ = -0.695, p < 0.01) displayed a strong negative correlation with age. The decline in stress again reached statistical significance between 26 and 68 weeks (from 304 ± 23 kPa to 188 ± 13 kPa; Figure 6B), while the decrease in stiffness was delayed to 84 weeks (from 1.74 ± 0.13 MPa to 1.10 ± 0.06 MPa; Figure 6B).

The distensibility of the wall decreased from 25.0 ± 1.8 MPa^-1^ at 12 weeks to 10.6 ± 0.8 MPa^-1^ at 84 weeks (Figures 1C & 6B) and was negatively correlated with age (r_s_ = -0.859, p < 0.01). The elastic energy responsible for blood flow augmentation also exhibited a negative correlation with age (r_s_ = -0.792, p < 0.01), whereby the first significant decline occurred between 26 and 68 weeks (90 ± 10 kPa vs. 48 ± 4 kPa; Figure 6B) and continued thereon to the last endpoint (38 ± 2 kPa at 84 weeks). These findings indicate that the structural stiffening of the ATA happens gradually throughout the mouse lifespan and that aging drastically impairs the ability of the ATA to passively deform and recoil under pulsatile pressure loads, which combined contribute to vascular dysfunction with advancing age.

### Effect of body mass on aortic diameter

Body mass imparted a significant effect on aortic caliber across age groups [F (6,43) = 6.770, p < 0.001]. Predicted systolic luminal diameters scaled allometrically rather than isometrically with body mass. Estimated values for the coefficients that describe the exponential relationship between the two metrics are *β* = 0.328 and *α* = 559 *m* · *g* ^*β*^ (R^2^ = 0.692; Figure 6D). Because of this relationship, the ratio between the luminal diameter of the normal ATA and body mass was not constant, but rather varied with body mass (Figure S3). Thus, normalization of aortic caliber by body mass should only be used as a comparison metric when no differences in body mass are observed across experimental groups^27^.

### Piezo-1-mediated enhanced aortic vasoconstriction with aging

To explore the molecular mechanisms underlying the age-related maladaptive remodeling of aortic tissues, we assessed the expression of the mechanosensitive ion channel Piezo-1 in the aged ATA. We have previously shown that Piezo-1 mediates pathological vascular remodeling during aortic aneurysms^28^. Piezo-1 expression was significantly increased in vSMCs ensconced in the media of the 84-week-old ATA (Figure 7A). α-actinin 2, a potent actin fiber crosslinker and the upstream regulator of Piezo-1, was also highly expressed in the ATA at 84 weeks, while aging did not affect α-smooth muscle actin expression of vSMCs in the media (Figure 7A). To determine the consequence of elevated Piezo-1 expression on the vasoconstrictor response of the aged ATA, we compared KCl-mediated contraction at 84 weeks, 12 weeks, and 12 weeks in the presence of the Piezo-1 activator Yoda1. Average active biaxial data revealed enhanced vasoconstriction with aging, under circumferentially isobaric (80mmHg transmural pressure) and axially isometric (vessel-specific *in vivo* stretch) conditions reproducing physiological loads. Following KCl administration, the decline in outer diameter occurred more rapidly in specimens from 84-week-old mice compared with the 12 weeks group (Figure 7B). As a result, by the end of the 15-minute testing window, the relative maximal contraction was larger in aged than young ATAs (16 ± 3% vs. 6 ± 1% relative decrease in outer diameter at 84 and 12 weeks, respectively; Figure 7C). Furthermore, activation of Piezo-1 in young ATAs following Yoda1 incubation supported average maximal contraction of 12 ± 1% relative to baseline outer diameter, thus recapitulating the enhanced vasoconstrictive response to KCl observed in aged samples (Figure 7C). Neither increasing age nor Yoda1 treatment altered the axial force during contraction (Figure 7B, 7C). Collectively, these results suggest that Piezo-1 could act as a critical gerogenic checkpoint by altering the structural/functional fate of the aorta during aging.

**Figure 7.**
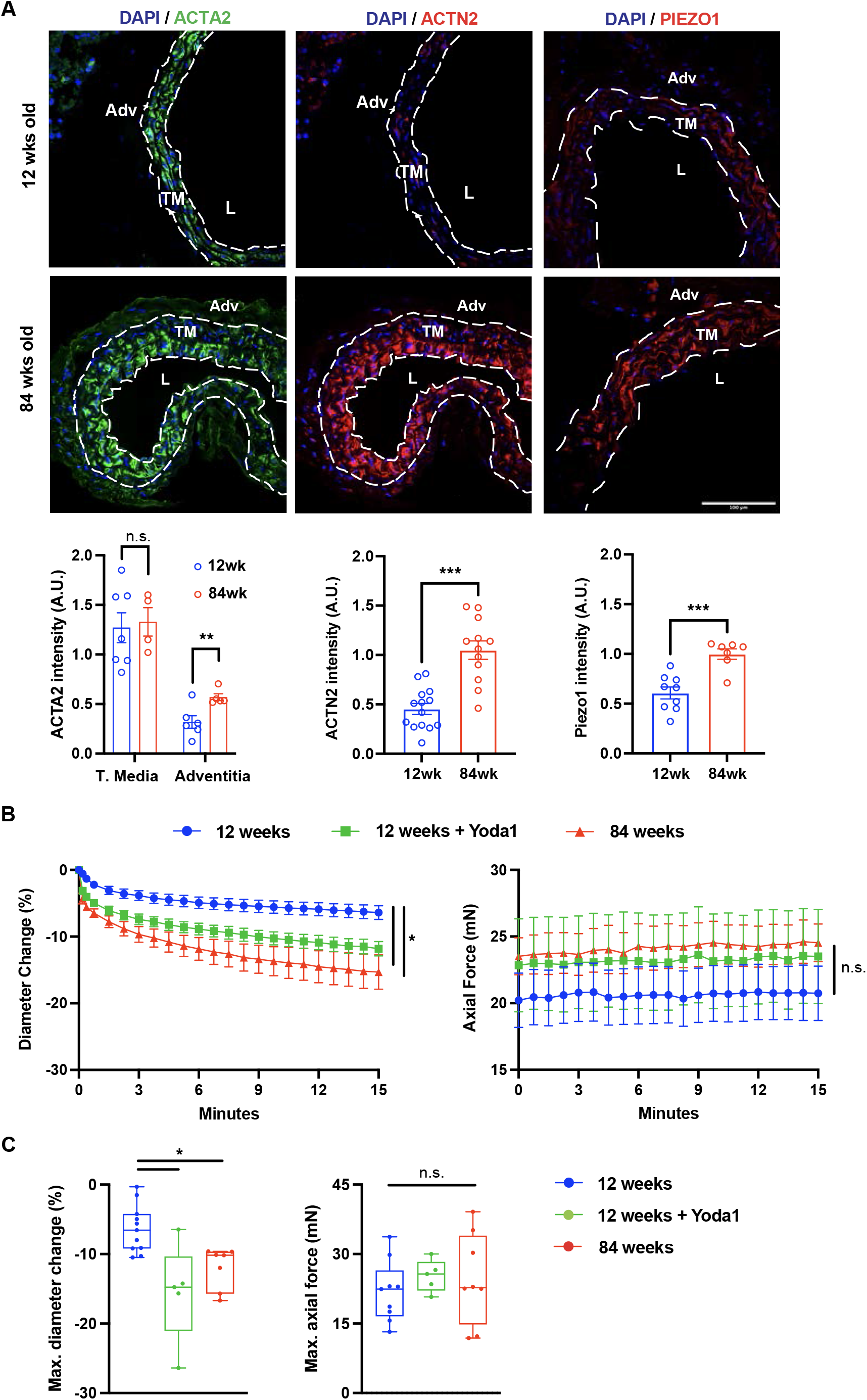
Piezo-1 activation enhances the vasoconstrictor response to KCl in the aged ATA. (A) Immunofluorescent staining colocalized with DAPI (cell nuclei, blue) for expression of smooth muscle α-2 actin (ACTA2, green), α-actinin 2 (ACTN2, red) and Piezo-1 (PIEZO1, red) in 84-week-old ATA tissues, compared to the 12-week endpoint. TM indicates the tunica media area outlined by white dotted lines, while Adv identifies the adventitial compartment, and L denotes the luminal edge of the vessel. Plots show relative fluorescent intensity, which was quantified separately in the tunica media and adventitia for ACTA2. Scale bar is 100μm. (B) Average (± SEM) traces of percent change in outer diameter and axial force following KCl stimulation of naïve ATA samples from 12-(N=11) and 84-week-old mice (N=5), and 12-week-old vessels preincubated with Yoda1 (N=7). (C) Maximum change in diameter and axial force in response to KCl for naïve ATA samples at the 12- and 84-weeks endpoint, as well as 12-week-old ATAs preincubated with Yoda1. Statistical significance between individual age groups denoted by ** overbar for p < 0.01 and *** overbar for p < 0.001.

## DISCUSSION

The consolidated relationship between vascular health and mortality^29^ corroborates the existence of a molecular dialogue between aging and cardiovascular disease^30,31^. Age-related changes in the structure and function of the heart and blood vessels alter the milieu in which pathophysiological processes operate^32^, influencing the threshold for disease inception, the rate of disease progression, and the severity of symptoms^33^. Likewise, pathological mechanisms may modify the natural course of the aging process, supporting a multi-organ accelerated decay. Motivated by our interest in the superposition between aging and environmental pollutant inhalation^27,34–36^, we probed cell makeup and gene/protein expression in tissues from the ascending thoracic aorta (ATA) of young and aged mice, to interpret the evolving geometrical features and functional metrics that this aortic segment exhibits throughout the mouse adult lifespan.

Alongside impairment of endothelium-dependent dilatation in resistance arteries^37–39^, structural stiffening of the aorta has emerged as a mechanical signature of aging across species, from mice^8,14,40–42^, rats^43^, and rabbits^44^ to nonhuman primates^45^ and humans^7,42^. We confirm here that distensibility declines in the murine ATA between 12 and 84 weeks of age (Figures 1C, 6B; Table S3) and further investigate the interplay between hemodynamics, geometry, and tissue stiffness through intermediate endpoints^33,40^. Besides wall thickness, both the luminal and outer diameters of the ATA progressively grow larger with age^8,40,41^ (Figure 6A; Table S3). Aortic caliber is primarily regulated by volumetric blood flow^46^ and varies allometrically with body mass^47^. While we did not measure cardiac output, the coefficients of allometric scaling^48,49^ that we estimated from predicted luminal diameters at physiological pressure (Figure 6D) show good agreement with those reported in previous studies^47,50,51^.

Progressive wall thickening and lumen widening of the ATA during aging rely primarily on collagen deposition and remodeling within the medial and adventitial layers of the wall (Figures 5D). While others have noted an overall increase of collagen in the aorta of aged mice^8,26,40,41^, Cavinato *et al*. focused on the first year of the mouse life and remarkably showed that collagen becomes more abundant and features thicker and less undulated fibers in the adventitia, while *de novo* collagen production in the media broadens the separation of adjacent elastic laminae^52^. The latter effect persists in the ATA of mice at 84 and 100^8^ weeks of age, reducing the relative content of medial elastin (Figure 5C)^8,26,40^. Despite apparent preserved integrity^8,26,40^ (Figure 5C), elastic fibers within central arteries of mice may also become less mechanically competent with age, due to increased enzymatic activity of elastase^26,53^. As a result of these layer-specific processes, mechanical loads are transferred to newly deposited/remodeled collagen fibers (or to stiffer smooth muscle bundles) and promote the circumferential stiffening of medial and adventitial tissues within the physiological range of loads^40^ (Figure 6B-C; Table S3). Compounded by the loss of axial extensibility, the layer-specific increase in circumferential stiffness of ATA tissues during aging impairs aortic function by reducing the elastic energy available for diastolic blood flow augmentation (Figure 6B; Table S3).

Substantial accumulation of T-lymphocytes in the aged ATA (Figure 4) upholds mounting evidence of immune system involvement in the adventitial stiffening of central arteries during aging^54,55^. Trott *et al*. recently showed that T-cell depletion significantly lowers aortic PWV and ameliorates mesenteric endothelium-dependent dilatation in ∼23-month-old male C57BL/6 mice^56^. Lesniewski *et al*. further related T-lymphocytes infiltration in perivascular and adventitial tissues with increased concentrations of pro-inflammatory cytokines in the thoracic aorta from male B6D2F1 mice aged ∼30 months^57^. Notably, Wu *et al*. reported that accumulation of activated T-cells enhances adventitial collagen deposition via cytokine release and stiffens thoracic aortic tissues from 6- and 9-month-old male mice with chronic vascular oxidative stress^58^, a prominent feature of aging^59,60^. Fleenor *et al*. further suggested that TGF-β1 overexpression in the adventitia of carotid arteries from ∼30-month-old B6D2F1 mice may promote collagen deposition and tissue stiffening by stimulating differentiation of fibroblasts into a synthetic phenotype expressing ACTA2^26^. Consonant with this speculation, ACTA2 expression in the adventitia is larger (Figure 7A) and fibroblasts from 84-week-old ATA tissues exhibit higher transcriptomic expression of collagen genes (Figure 5B). Therefore, T-lymphocyte infiltration (Figure 4M) may contribute to adventitial tissue stiffening (Figure 6B-C) by promoting fibrosis (Figure 5D) through activation of fibroblasts (Figures 5B, 7A) in the aging ATA.

Lack of T-cell accumulation substantiates medial immune-privilege^61^ and shifts the mechanistic burden for the age-related stiffening of this aortic compartment onto vSMCs. Depletion of vSMCs in the 84-week-old ATA (Figure 2E-G) corroborates histological evidence showing progressive decrease in their area fraction during aging^8,41^ (Figure S2B). Nevertheless, preserved smooth muscle α-2 actin (ACTA2) fluorescence intensity (Figure 7A) in the media alludes to the retention of a contractile phenotype by aged vSMCs. Transcriptional upregulation of netrin signaling (Figure 3A) at the 84-week endpoint prompted evaluation of Piezo-1 activity as a candidate mediator of age-related remodeling in the ATA media. We have recently shown that actin fiber crosslinking and cytoskeletal stiffening due to netrin-induced expression of α-actinin 2 mediate Piezo-1 opening in vSMCs, thereby promoting Ca^2+^ influx and proteolytic enzyme release in the aneurysmal abdominal aorta^28^. Tissue expressions of α-actinin 2 (ACTN2) and Piezo-1 (PIEZO1) significantly intensify in the ATA media of aged mice (Figure 7A), while augmented vasoconstriction of ATA specimens under physiological biaxial loads^62–65^ suggests Piezo-1 activation at the 84-week endpoint (Figure 7B-C). In support of the latter, recapitulation of α-actinin 2-mediated powering of Piezo-1 through Yoda1 abatement of the mechanical threshold for channel engagement^66^ also enhances vasoconstriction of 12-week-old ATAs (Figure 7B-C). Aligned with our findings, Liao *et al*. reported that Piezo-1 activation by Yoda1 induces contraction of human pulmonary arterial vSMCs and dose-dependent vasoconstriction of rat intrapulmonary arteries, due to intracellular Ca^2+^ increase^67^. These observations fuel maturing evidence of augmented calcium signaling^68^ and contractility^69,70^ in SMCs embedded within a stiff matrix. Note, Retailleau *et al*. reported that Piezo-1 signaling promotes thickening of the murine caudal artery by stimulating the Ca^2+^-dependent catalytic activity of transglutaminases^71^ and tissue transglutaminase contributes to vascular stiffening during aging^72–74^. Therefore, modulation of the actin cytoskeletal architecture in vSMC (Figure 7A) may heighten contractility (Figure 7B-C) and contribute to ECM stiffening (Figure 6B-C; Table S3) by mediating expression and activation of mechanosensitive Piezo-1 in the ATA media during aging (Figure 7A).

In conclusion, we have shown that chronological aging of the aorta is a complex process involving layer-specific multicellular and tissue micro-adaptations that manifest as a region-dependent and progressive loss of function. We demonstrated that the structural stiffening is heterogeneous along the aortic tree, with the ATA exhibiting the most stringent decline in distensibility. We also discussed two separate molecular mechanisms that may independently support the collagen-driven stiffening of the medial and adventitial layers of the wall, respectively. While T-lymphocyte regulation of adventitial fibroblasts likely upholds centrifugal fibrosis, the α-actinin 2/Piezo-1 axis rather mediates a centripetal aging of medial vSMCs that may promote tissue remodeling. These new results further refine our understanding of the mechanisms that advance chronological vascular aging and provide a candidate platform of signals that could be investigated in ailments affecting the aorta.

## MATERIALS AND METHODS

### Animals and physiological measurements

All animal procedures adhered to NIH guidelines and received approval from the Institutional Animal Care and Use Committee (IACUC) at Northeastern University or New York University School of Medicine. Female C57BL/6 mice were purchased from Jackson Laboratory (Bar Harbor, ME) at 7-8 weeks of age and housed in groups of 5 under standard 12:12-h light-dark cycle, with *ad libitum* access to water and chow. Mice were allowed to age up to either 12±1 weeks (N=25), 26±1 weeks (N=8), 49±2 weeks (N=8), 68±1 weeks (N=9), or 84±4 weeks (N=32) by random assignment. These endpoints cover most of the mouse adult lifespan and precede the stark age-related decline in survival beyond ∼100 weeks of age^75^ (Figure S1). Body mass and tail-cuff peripheral pressure (CODA, Kent Scientific) were measured in a subset of mice at endpoint, before CO2 euthanasia.

### Single-cell RNA sequencing

Ascending thoracic aorta (ATA) sections were obtained from young (12±1 week-old, N=3) and aged (84±4 week-old, N=3) mice for single-cell RNA sequencing (scRNA-seq). Tissues were digested for 60 minutes at 37°C in an enzymatic mix composed of type II collagenase (10 mg/ml; C6885, Sigma Aldrich) and elastase (1 mg/ml; LS002292, Worthington Biochemistry), as previously described^24,28^. Single cell suspensions were obtained after passage of the digested lysate through a 70 μm cell-strainer. Following digestion, viable cells were enriched using a dead cell removal magnetic kit (130-090-101, Miltenyi Biotech) and cellular viability was assessed using Trypan blue staining (0.4%; 1450013, Bio-Rad) and an automated cell counter (TC20, Bio-Rad). Viable cells (15×10^3^ per sample) were loaded on a 10x Genomics Chromium instrument to obtain individual gel beads in emulsion (GEMs). Library constructions were prepared using Chromium Single Cell 3′Reagent Kits v.2 (10x Genomics; PN-120237, PN-120236, PN-120262).

The HiSeq 4000 (Illumina) was used for sequencing with 2 × 150 paired-end reads (>90% sequencing saturation). Cell Ranger Single Cell Software Suite (version 1.3) was used to perform de-multiplexing, barcode and UMI processing, and single-cell 3′ gene counting (https://support.10xgenomics.com/single-cell-gene). Data analysis was performed using the Seurat sequencing package (version 4.0.5), R Studio Desktop (version 1.2.5033), and R (version 3.0.1+). Quality control, metrics, data normalization, scaling, batch correction, and dimension reductions were all performed using the Seurat package. Cells with a mitochondrial transcript proportion of <5% were kept for analysis, while neighboring and clustering was performed on the most significant principal component analysis.

### Active biaxial mechanical testing

Immediately after sacrifice of young (12±1 week-old, N=11) and aged (84±4 week-old, N=5) mice, ascending thoracic aorta (ATA) specimens were excised and immersed in Hanks buffered salt solution (HBSS) chilled to 4°C. Excess perivascular tissue was quickly removed and lateral branches were ligated using one of three strands of a braided 9-0 nylon suture. Vessels were secured onto a stage using custom glass cannulae and immersed in a 37°C Krebs–Ringer’s buffer solution, bubbled with a 95% O_2_ – 5% CO_2_ mixture to maintain pH at 7.4. The stage was mounted onto a computer-controlled custom biaxial tester that applies luminal pressure and axial stretch while measuring outer diameter and axial force. Following preconditioning (40mmHg pressure at 1.1 axial stretch + 60mmHg pressure at 1.2 axial stretch), refinement of initial measurements of unloaded dimensions facilitated estimation of the *in vivo* axial stretch. Vessels were exposed to an 80mmHg distending pressure while maintained at *in vivo* axial stretch and allowed to equilibrate. A subset of 12-week-old specimens (N=7) was incubated for 30 minutes with 20μM Yoda1 to activate Piezo-1 mechanosensitive channels. Upon stabilization of outer diameter and axial force, vasoconstriction was induced by addition of a 40mM KCl to the bath, which was washed out with fresh Krebs-Ringer’s solution 15 minutes later. Vasoconstriction throughout was expressed as a percent change in outer diameter,

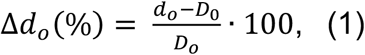

where *d*_0_ is the current outer diameter and *D*_0_ is the initial outer diameter prior to KCl treatment.

### Passive biaxial mechanical testing

The custom biaxial device used for probing KCl vasoconstriction also facilitated passive mechanical testing, according to experimental protocols and analytical methods described elsewhere^27,76,77^. Briefly, following dissection, removal of perivascular tissue, and ligation of lateral branches, each aorta was sectioned into ascending (ATA, N=43) and descending (DTA, N=50) thoracic and suprarenal (SAA, N=10) and infrarenal (IAA, N=10) abdominal segments. Cannulated specimens were axially stretched and preconditioned under physiological pulsatile loads while submerged in room temperature HBSS to ensure a near passive mechanical behavior^78^. Upon refinement of preliminary unloaded dimension and *in vivo* axial stretch estimates, the pressure-diameter (at varying axial stretch) and force-length (at varying pressure) responses were recorded through a set of 7 cyclic testing protocols.

### Transmurally-averaged mechanical properties

Experimental data were supplied to a nonlinear regression algorithm to fit a microstructurally motivated four-fiber family strain energy potential (*W*) in the form

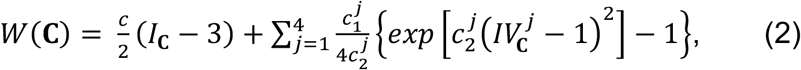

where *I*_*C*_ and 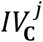 are the first and fourth invariants of the right Cauchy-Green deformation tensor **C** and index *j* encompasses the axial (*j* = 1), circumferential (*j* = 2), and two axially symmetric diagonal (*j* = 3,4) directions. The first isotropic Neo-Hookean term (coefficient *c*, dimension of a stress) in Equation 2 accounts for the contribution of elastic fibers and amorphous matrix, while the Fung-type terms (coefficients 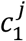 with the dimension of a stress and unitless coefficients 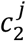) describe the anisotropic response of collagen and smooth muscle bundles^79^. The 7 best-fit parameters were used to predict geometry and transmurally-averaged tissue properties under desired pressure and axial loads, leveraging the small-on-large theory to determine the circumferential and axial components of linearized stiffness about the set working point within the cardiac cycle^80^. Based on these predictions, the structural stiffness metric of cyclic aortic distensibility (*𝒟*) was calculated from the systolic and diastolic values of blood pressure (*P*) and inner diameter (*d*_*i*_), as

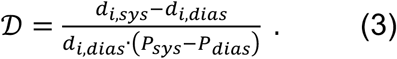

### Layer-specific tissue stiffness

Biaxial data were further analyzed using a bi-layered model of the aortic wall that features microstructurally-motivated and layer-specific constitutive relations and accounts for the deposition stretches of newly secreted constituents (e.g., elastic fibers or multiple families of collagen fibers) as they integrate within the existing matrix^76^ in the homeostatic reference configuration. We assumed a mass-averaged strain energy function in the form:

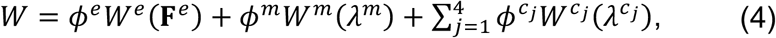

where the superscripts *i* = *e, m* and *c* refer to elastic fibers, smooth muscle bundles, and each of four families of collagen fibers (*j* = 1,2,3,4), respectively, *ϕ*^*i*^ are the mass fractions (from histological analysis, Figures 5 and S2) and *W*^*i*^ are the stored energy functions for the constituents that compose the mixture, **F**^*e*^ is the deformation gradient tensor experienced by the elastic fibers, and *λ*^*m*^ and 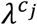 are the stretches experienced by the smooth muscle and the *j*^*th*^ family of collagen fibers, respectively. Similar to the bulk formulation in Equation 2, we described the elastic fiber behavior with a neo-Hookean stored energy function

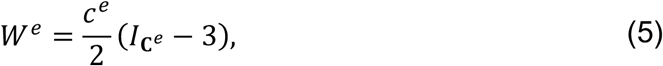

Where *c*^*e*^ is a coefficient with the dimension of a stress, 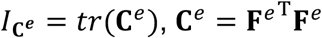 and 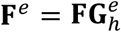, with **F** the deformation gradient tensor that describes the deformation of the mixture as a whole and 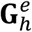 the deposition stretch tensor between the natural (stress-free) configuration of the elastic fibers and the homeostatic reference configuration. 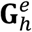 was assumed to be diagonal with circumferential and axial components within the [1.65, 1.74] range^77^ and the radial stretch determined by enforcing incompressibility. The nonlinear responses of collagen fibers and circumferential smooth muscle were modeled as Fung exponentials,

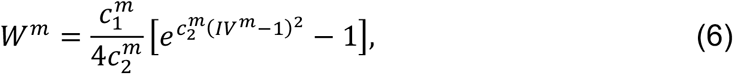

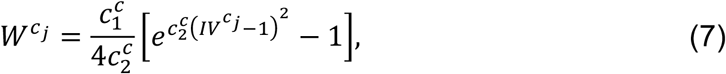

Where 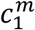 and 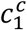 are coefficients with the dimension of a stress, while 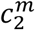 and 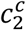 are dimensionless. Neither smooth muscle nor collagen fibers were assumed to have any radial component. The stretch experienced by smooth muscle was obtained by projecting ***C*** along the cell axis,

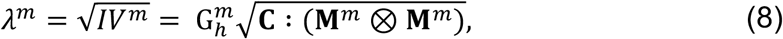

Where 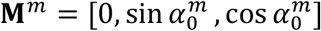 is the reference smooth muscle orientation and 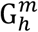 assumed values in the [1.03, 1.05] range^77^. Similarly, the stretch in the collagen fiber direction was determined as

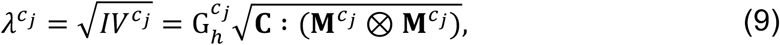

Where 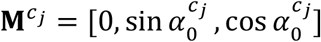 is the reference dominant orientation of the *j*^*th*^ collagen fiber family and 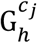 varied within the [1.10, 1.13] range^77^. Also estimated from experimental data were two additional parameters for the proportion of circumferentially-(*β*_*ϑ*_) and axially-(*β*_*z*_) oriented collagen fibers. The 8 best-fit parameters were obtained via nonlinear regression on the average experimental data and used to predict the transmural distribution of linearized circumferential tissue stiffness following the small-on-large approach^80^.

### Allometric scaling

Logarithmic transformation and linear regression of estimated systolic dimensions were used to yield best fit values for the *α* and *β* coefficients of the allometric scaling

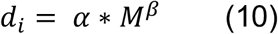

that relates luminal aortic diameter (*d*_*i*_) to body mass (*M*).

### Histology and Immunofluorescence

Following completion of mechanical testing, ATA tissues reserved for histology were fixed overnight in 10% neutral buffered formalin and stored long-term in 70% ethanol. Paraffin-embedded samples were serially sliced into 5μm thick cross sections that started ∼500mm distally of the aortic root. Once deparaffinized, cross sections were processed for tissue rehydration and antigen retrieval using EDTA and Tween solution. Tissues were stained with Verhoeff Van Gieson (VVG) or Masson’s Trichrome (MTC) stains. Images of stained cross sections were acquired individually with a 20x objective. The area fractions of elastin and collagen were quantified by automatic thresholding of pixels based on hue (H, 0-360), saturation (S, 0-1), and lightness (L, 0-1) values specific to each stain^27,81^. Following removal of cell nuclei and debris, all pixels with lightness below 0.30 in VVG-stained tissues were counted as elastin. Likewise, blue pixels (H 160-265, S 0.23-1.00, L 0.38-1.00) were considered as collagen in MTC-stained cross sections.

Additional ATA samples were dissected and immediately embedded in optimal cutting temperature (OCT) compound (4585, Fisher Scientific) and fast-frozen for cryosectioning. Tissues were sliced into 7μm thick cross-sections and staining was performed by overnight incubation with primary antibodies: anti-CD3 (#ab467053, Invitrogen, 1:100 dilution), anti-*α*-actinin2 (#14221-1-AP, Proteintech, 1:200 dilution), anti-actin-*α*-2 (#48938S, Cell Signaling, 1:200 dilution), anti-Piezo1, (#15939-1-AP, Proteintech and #APC-087, Alomone labs, 1:200 dilution each), and anti-HMGB1 (#ab11354, Abcam, 1:200 dilution). Alexa Fluor 488 and 568 conjugated anti-IgG antibodies (Invitrogen) were used for fluorescent signal detection (1:500 dilution each). 4’,6-diamidino-2-phenylindole (DAPI; Invitrogen) allowed visualization of the nucleus (1:5000 dilution). Images were acquired at x40 magnification using a Zeiss LSM 710 confocal microscope (Carl Zeiss) and the Zeiss Efficient Navigation (ZEN) software (Carl Zeiss). Identical acquisition parameters were set to capture images from 12-week and 84-week age groups. Mean fluorescent intensity per unit area was computed using ImageJ software (NIH).

### Statistics

Unless otherwise noted, all values are reported as average ± SEM. Due to lack of normality, the one-way non-parametric Kruskal-Wallis test with Bonferroni correction for multiple comparisons was chosen to evaluate differences in geometrical, mechanical, and microstructural metrics across age groups. The non-parametric Spearman’s rank-order correlation coefficient was used to further measure the strength and direction of association between increasing age and metrics of interest. Significance difference in vasoconstriction parameters was evaluated using one-way ANOVA. The effect of age, body mass, and their interaction on aortic geometry was assessed using one-way ANCOVA with age group as the independent variable and body mass as the covariate. For scRNA-seq data, p-values were adjusted using the Benjamini-Hochberg method for false discovery correction. Genes with an adjusted p-value < 0.05 were considered as differentially expressed.

## DATA AVAILABILITY

The authors declare that all data supporting the results in this study are reported within the Manuscript and its Supplementary Information. Raw data are available from the corresponding authors on reasonable request.

## ACKNOWLEDGMENTS

We acknowledge financial support from the National Institute of Health (R01 HL146627 and R01HL149927 to B.R. and R21 HL148747 to C.B). M.S is funded by American Heart Association post-doctoral fellowship (907602).

## AUTHOR CONTRIBUTIONS

CB and BR conceptually developed and supervised all aspects of the project. YF and JM performed biomechanical studies and data analysis. MS and CM performed sequencing and analysis. YF, CB, and BR wrote the manuscript assisted by CM. YP, PK and JV performed experiments and helped with graphical illustration.

