## Supplementary Material for "Mapping the unicellular transcriptome of the ascending thoracic aorta to changes in mechanosensing and mechanoadaptation during aging"

#### FIGURES & TABLES

**Table S1**. Average (± SEM) age, body mass, and blood pressure across age groups.

| | Body mass (g) | Blood Pressure (mmHg) | | Age | | |
|---|---|---|---|---|---|---|
| | | Systolic | Diastolic | Days | Weeks | Months |
| *12* | 19.9 ± 0.4 | 113 ± 6 | 81 ± 6 | 86 ± 1 | 12.3 ± 0.2 | 2.83 ± 0.05 |
| *26* | 23.2 ± 0.8 | 118 ± 5 | 88 ± 2 | 179 ± 2 | 25.5 ± 0.3 | 5.87 ± 0.07 |
| *49* | 26.5 ± 1.0 | 112 ± 3 | 76 ± 3 | 343 ± 4 | 48.9 ± 0.5 | 11.25 ± 0.12 |
| *68* | 30.7 ± 1.3 | 106 ± 1 | 76 ± 2 | 476 ± 1 | 68.0 ± 0.2 | 15.65 ± 0.05 |
| *84* | 30.0 ± 0.7 | 107 ± 3 | 70 ± 7 | 590 ± 6 | 84.3 ± 0.8 | 19.40 ± 0.18 |

**Table S2**. Best-fit coefficients of the four-fiber family strain energy potential (Equation 2) as estimated from biaxial experimental data to describe the bulk mechanical response of the ascending thoracic aorta (ATA) at the 12-, 26-, 49-, 68-, and 84-week endpoints.

| | Elastic Fibers | Axial Collagen | | Circumferential Collagen + SMC | | Symmetric Diagonal Collagen | | | Error |
|---|---|---|---|---|---|---|---|---|---|
| | $c\ (kPa)$ | $c_1^1\ (kPa)$ | $c_2^1$ | $c_1^2\ (kPa)$ | $c_2^2$ | $c_1^{3,4}\ (kPa)$ | $c_1^{3,4}$ | $\alpha_0$ (deg) | $RMSE$ |
| **ATA** | | | | | | | | | |
| *12 week* | 19.151 | 13.540 | 2.3E- | 16.198 | 8.2E-13 | 14.442 | 0.374 | 46.043 | 0.052 |
| *26 week* | 26.647 | 8.299 | 2.3E- | 18.411 | 0.068 | 10.848 | 0.3915 | 47.0135 | 0.051 |
| *49 week* | 8.845 | 13.792 | 5.3E- | 4.6E-08 | 6.721 | 18.439 | 0.291 | 52.677 | 0.061 |
| *68 week* | 6.319 | 17.764 | 2.8E- | 1.0E-04 | 5.005 | 26.953 | 0.359 | 51.451 | 0.065 |
| *84 week* | 7.106 | 18.614 | 0.017 | 3.1E-05 | 6.526 | 23.878 | 0.538 | 51.589 | 0.051 |

**Table S3.** Morphological and mechanical properties of the ascending thoracic aorta (ATA) at the 12-, 26-, 49-, 68-, and 84-week endpoints. Statistical significance for the difference between age groups is denoted by * for $p < 0.05$ *vs.* 12 weeks, † for $p < 0.05$ *vs.* 26 weeks, ‡ for $p < 0.05$ *vs.* 49 weeks, and § for $p < 0.05$ *vs.* 68 weeks. Spearman correlation coefficients ($r_s$) that relate mechanical metrics to age are also reported, with significance indicated by ¥ for $p < 0.05$.

| ATA | 12 weeks | 26 weeks | 49 weeks | 68 weeks | | 84 weeks | | Spearman | |
|---|---|---|---|---|---|---|---|---|---|
| n | 6 | 8 | 7 | 8 | | 14 | | $r_s$ | |
| Distensibility ($MPa^{-1}$) | 25.0 ± 1.8 | 24.7 ± 1.6 | 15.4 ± 0.5 | 15.8 ± 1.0 | | 10.6 ± 0.8 | *† | -0.859 | ¥ |
| ***Unloaded*** | | | | | | | | | |
| Outer Diameter (µm) | 1115 ± 26 | 1109 ± 32 | 1186 ± 37 | 1176 ± 27 | | 1353 ± 25 | *†§ | 0.752 | ¥ |
| Wall Thickness (µm) | 111 ± 1 | 108 ± 6 | 128 ± 2 | 128 ± 2 | | 146 ± 5 | *† | 0.808 | ¥ |
| ***Loaded systolic pressure*** | | | | | | | | | |
| Outer Diameter (µm) | 1639 ± 24 | 1642 ± 15 | 1737 ± 42 | 1646 ± 64 | | 1912 ± 23 | *†§ | 0.688 | ¥ |
| Wall Thickness (µm) | 40 ± 1 | 40 ± 4 | 51 ± 3 | 54 ± 1 | | 65 ± 2 | *† | 0.759 | ¥ |
| Stretch (-) | | | | | | | | | |
| Circumferential | 1.60 ± 0.03 | 1.61 ± 0.05 | 1.62 ± 0.04 | 1.57 ± 0.03 | | 1.53 ± 0.02 | † | -0.275 | |
| Axial | 1.76 ± 0.04 | 1.72 ± 0.04 | 1.68 ± 0.05 | 1.51 ± 0.02 | † | 1.47 ± 0.03 | *†‡ | -0.738 | ¥ |
| Cauchy Stress (kPa) | | | | | | | | | |
| Circumferential | 315 ± 11 | 331 ± 27 | 261 ± 15 | 160 ± 7 | † | 222 ± 7 | † | -0.574 | ¥ |
| Axial | 316 ± 14 | 304 ± 23 | 242 ± 11 | 188 ± 13 | † | 177 ± 9 | *† | -0.707 | ¥ |
| Linearized Stiffness (MPa) | | | | | | | | | |
| Circumferential | 1.98 ± 0.14 | 2.24 ± 0.23 | 2.66 ± 0.21 | 2.55 ± 0.42 | | 3.73 ± 0.40 | * | 0.521 | ¥ |
| Axial | 1.78 ± 0.11 | 1.74 ± 0.13 | 1.42 ± 0.06 | 1.33 ± 0.11 | | 1.10 ± 0.06 | *† | -0.695 | ¥ |
| Stored Energy (kPa) | 88 ± 3 | 90 ± 10 | 61 ± 4 | 48 ± 4 | † | 38 ± 2 | *†‡ | -0.792 | ¥ |

**Table S4**. Best-fit parameters of a mass-averaged form of the four-fiber family strain energy potential (Equations 4-7) as estimated from biaxial experimental data to describe the layer-specific mechanical response of the ascending thoracic aorta (ATA) at the 12- and 84- week endpoints.

| | Elastic Fibers | SMCs | | Collagen Fibers | | | | | Error |
|---|---|---|---|---|---|---|---|---|---|
| | $c^e$ $(kPa)$ | $c_1^m$ $(kPa)$ | $c_2^m$ | $c_1^c$ $(kPa)$ | $c_2^c$ | $\alpha_0$ (deg) | $\beta_\vartheta$ | $\beta_z$ | $RMSE$ |
| **ATA** | | | | | | | | | |
| *12 week* | 179.660 | 471.226 | 4.711 | 4389.129 | 1.232 | 45.948 | 0.072 | 0.031 | 0.052 |
| *84 week* | 266.285 | 497.360 | 60.069 | 1279.611 | 8.379 | 46.971 | 0.011 | 0.014 | 0.054 |

**Figure Captions**

**Figure S1.** Survival curve for female C57BL/6 mice, recreated with data from the Jackson Laboratory Yuan2 dataset (RRID: SCR_003212). The red shaded area envelopes the 95% confidence interval. The endpoints for this study are visualized by vertical bars and cover most of the mouse adult lifespan, before the probability of survival begins to decline after ~100 weeks of age.

**Figure S2.** Collagen remodeling supports the microstructural aging of the ATA wall. The relative collagen content progressively increases with age in the ATA wall (A), while the area fraction of elastin and vSMC both decline (B). Statistical significance between individual age groups denoted by * overbar for $p < 0.05$.

**Figure S3.** The expected ratio between aortic caliber and body mass varies with body mass, emphasizing that normalization by body mass should be used to compare diameters across mice of similar size.

### Figure S1

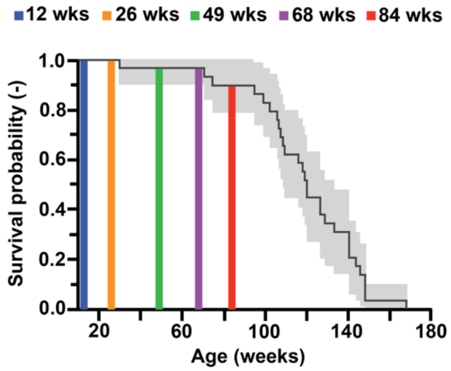
12 wks
26 wks
49 wks
68 wks
84 wks
1.0
0.8
0.6
0.4
0.2
0.0
Survival probability (-)
20
60
100
140
180
Age (weeks)

#### Figure S2

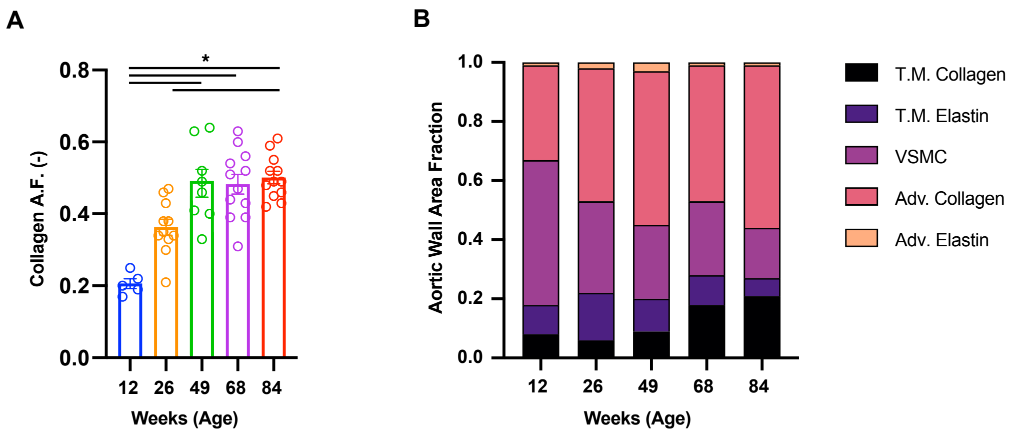
A
B
Collagen A.F. (-)
Weeks (Age)
Aortic Wall Area Fraction
Weeks (Age)
T.M. Collagen
T.M. Elastin
VSMC
Adv. Collagen
Adv. Elastin

### Figure S3

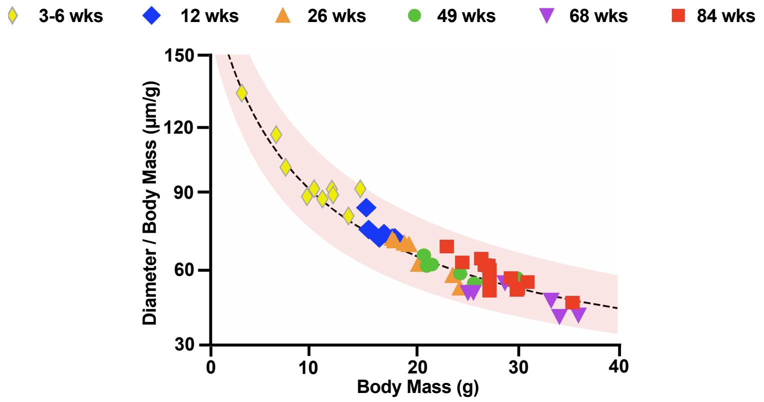

3-6 wks
12 wks
26 wks
49 wks
68 wks
84 wks
Diameter / Body Mass (µm/g)
150
120
90
60
30
0
10
20
30
40
Body Mass (g)
